# Filament structures unveil the dynamic organization of human acetyl-CoA carboxylase

**DOI:** 10.1101/2024.06.16.599241

**Authors:** Fayang Zhou, Yuanyuan Zhang, Yuyao Zhu, Qiang Zhou, Yigong Shi, Qi Hu

**Affiliations:** China College of Life Sciences, Zhejiang University; Hangzhou 310058, China; School of Life Sciences, Westlake University; Hangzhou 310024, China; Zhejiang Key Laboratory of Structural Biology, School of Life Sciences, Westlake University; Hangzhou 310024, China; Westlake AI Therapeutics Lab, Westlake Laboratory of Life Sciences and Biomedicine; Hangzhou 310058, China

## Abstract

Human acetyl-CoA carboxylases (ACCs) catalyze the carboxylation of acetyl-CoA, which is the rate-limiting step in fatty acid synthesis. The molecular mechanism underlying the dynamic organization of ACCs is largely unknown. Here, we determined the cryo-EM structure of human ACC1 in its inactive state, which forms a unique filament structure and is in complex with acetyl-CoA. We also determined the cryo-EM structure of human ACC1 activated by dephosphorylation and citrate treatment, at a resolution of 2.55 Å. Notably, the covalently linked biotin binds to a site that is distant from the acetyl-CoA binding site when acetyl-CoA is absent, suggesting a potential coordination between biotin binding and acetyl-CoA binding. These findings provide insights into the structural dynamics and regulatory mechanisms of human ACCs.

**Teaser:** Cryo-EM structures reveal how human acetyl-CoA carboxylase 1 changes shape to control its activity.

## Introduction

Acetyl-CoA carboxylases (ACCs) in the biotin-dependent carboxylase (BDC) family catalyze the reaction between acetyl-CoA and bicarbonate to produce malonyl-CoA(*1*). This reaction, known as the carboxylation of acetyl-CoA, is the first and rate-limiting step in fatty acid synthesis, thereby playing a crucial role in lipid metabolism(*2*).

ACCs exist ubiquitously in organisms ranging from prokaryotes to eukaryotes. Prokaryotic ACCs are multi-subunit complexes that consist of the biotin carboxyl carrier protein (BCCP), biotin carboxylase (BC), and carboxyltransferase (CT) subunits(*3*). The BCCP subunit is firstly biotinylated at the lysine residue in its A/VMKM motif with the catalysis of biotin-protein ligase (BPL)(*4*). Then, the BCCP subunit binds to the BC subunit, where the covalently bound biotin is carboxylated using bicarbonate as the carboxyl group donor and ATP as an energy source(*5, 6*). Lastly, the BCCP subunit relocates to the CT subunit, where the carboxylated biotin transfers its carboxyl group to acetyl-CoA to complete the entire catalytic process(*5, 6*). Unlike prokaryotic ACCs, eukaryotic ACCs are single-chain proteins with the BC, BCCP, and CT domains arranged sequentially(*3, 6*). Additionally, there is a BC-CT interaction (BT) domain between the BC and BCCP domains and a central domain (CD) between the BCCP and CT domains in each eukaryotic ACC(*6*).

Crystal structures of the BC subunit of *Escherichia coli* ACC, as a representative of prokaryotic ACCs, reveal a homodimeric organization of the BC subunit and explain its catalytic mechanism(*7*). In contrast, eukaryotic BC domains alone are inactive monomers(*8*). A crystal structure of *Saccharomyces cerevisiae* ACC (*Sc*ACC), in which *Sc*ACC forms a homodimer, shows that the BC domain is activated through dimerization(*8*). In addition to *Sc*ACC, which is expressed in the cytosol, *S. cerevisiae* has a second acetyl-CoA carboxylase called Hfa1p, which is expressed in the mitochondria(*9*).

There are also two isoforms of ACC in humans: ACC1, expressed in the cytosol, and ACC2, associated with the outer membrane of mitochondria (*10*). Human ACC1 and ACC2 share a sequence identity of 75%(*6*). Their catalytic activities are regulated by phosphorylation and direct binding of small molecule or protein regulators(*11-14*).

Phosphorylation leads to the inactivation of human ACCs(*12*). Citrate, a key metabolite in the citric acid cycle, activates human ACCs by binding to them and regulating the organization of different domains within the enzymes(*15, 16*). Dephosphorylation and citrate binding have an additive effect in activating human ACCs(*12,17,18*). In the presence of citrate, the dephosphorylated ACC1 forms filaments (ACC-citrate filaments)(*16*). A cryo-EM structure of the ACC-citrate filaments, at a resolution of 5.4 Å, reveals that the filaments are formed by the assembly of ACC1 homodimers mediated by the CD domain(*16*). The structure of the ACC1 homodimer in the ACC-citrate filament is similar to that of the *Sc*ACC homodimer, suggesting a conserved working mechanism of eukaryotic ACCs(*8*, *16*).

In contrast to the activating effect of citrate, the breast cancer type 1 susceptibility protein (BRCA1) recognizes the phosphorylated human ACC1 through its BRCT domain to inhibit the catalytic activity of ACC1(*19*). The binding of the BRCT domain to ACC1 results in the formation of inactive ACC1 filaments (ACC-BRCT filaments)(*16*). A cryo-EM structure of the ACC-BRCT filaments shows that the BC domain is monomeric, and the BCCP domain cannot reach the active site of the BC domain, explaining why ACC1 is inactive in the ACC-BRCT filaments(*16*).

In this study, we purified endogenously expressed human ACC from Expi 293F cells and identified a new type of ACC filaments (ACC-ACO filaments, in which the term “ACO” refers to the acetyl-CoA-bound state of ACC) by incubating the purified ACC with the substrates and determining the cryo-EM structure. We also determined a cryo-EM structure of the ACC-citrate filaments at an improved resolution. Structure analysis indicates that ACC1 in the ACC-ACO filaments adopts an inactive conformation. Our structures, together with the cryo-EM structures of the ACC-citrate filaments and the ACC-BRCT filaments reported previously, demonstrates that the organization of human ACCs is highly dynamic, with the dynamics being regulated by various regulators.

## Results

### Preparation of the human ACC1 samples for cryo-EM study

Using an engineered streptavidin immobilized on agarose resin (the Strep-Tactin^®^XT resin), we extracted human biotin-dependent carboxylases (BDCs) endogenously expressed in Expi 293F cells. The engineered streptavidin recognizes the biotin that is covalently linked to the BCCP domains of BDCs. Therefore, the BDCs can be extracted without the need for adding any affinity tags. The extracted BDCs were further purified by size exclusion chromatography and analyzed by SDS-PAGE (fig. S1A–C, E–G). Each band on the SDS-PAGE gel was analyzed separately using mass spectrometry. For the top band, ACC1 and ACC2 ranked first and second on the protein list identified by mass spectrometry (table S1). However, only one unique peptide fragment corresponding to ACC2 was identified, suggesting that the top band corresponds to ACC1.

The other three types of human BDCs, including pyruvate carboxylase (PYC), propionyl-CoA carboxylase (PCC), and methylcrotonyl-CoA carboxylase (MCC), were also identified (fig. S1C, G).

To prepare the ACC-citrate filaments, the fractions with elution volumes between 12 to 13.5 mL, in which ACC1 was a major component, were pooled, dephosphorylated by protein phosphatase PP1α, and treated with citrate (fig. S1A–D). To prepare ACC in complex with its substrates, the fractions from another batch of BDCs with elution volumes between 12.5 and 16.5 mL were pooled and incubated with acetyl-CoA, bicarbonate, MgCl_2_, and ATP (fig. S1E–H).

### Overall structures of the human ACC filaments

The structures of the ACC-citrate filaments and ACC in complex with its substrates were determined using cryo-EM, resulting in overall resolutions of 4.14 Å and 2.73 Å, respectively (figs. S2, S3 and table S2). ACC in complex with its substrates also forms filaments and is referred to as ACC-ACO filaments. Further data processing, focusing on the core regions of the two types of filaments which mainly contain an ACC homodimer, yielded structures at improved overall resolutions of 2.55 Å (ACC-citrate filaments) and 2.57 Å (ACC-ACO filaments) (figs. S2, S3 and table S2). In the ACC-ACO filaments, the cryo-EM density of each BC domain is only partially visible (fig. S4A). Nevertheless, by considering the position of the BT domain, we were able to dock the BC domain into the cryo-EM map (fig. S4B). We will focus on the 2.55 Å and 2.57 Å structures in the following analysis.

Since there are two isoforms of ACC (ACC1 and ACC2) in human cells, we docked ACC1 and ACC2 separately into the cryo-EM map of the core region of the ACC-citrate filaments (fig. S5A–C) to determine which isoform the cryo-EM structure corresponds to. We focused on the region comprising residues 1601–1640 in ACC1 (corresponding to residues 1720– 1751 in ACC2). In this region, ACC1 has 8 additional residues (fig. S5D). The structure of ACC1 aligns better with the cryo-EM map. Similar analysis indicates that the cryo-EM map of the core region of the ACC-ACO filaments also matches better with ACC1. These findings are consistent with the mass spectrometry data (table S1). Therefore, we built the structure models for both the ACC-citrate filaments and the ACC-ACO filaments based on the sequence of ACC1.

The overall structure of the ACC-citrate filaments closely resembles the previously reported 5.4 Å structure (PDB code: 6G2D)(*16*). But because of the higher resolution of our structure, each domain of ACC1 is clearly defined, and the side chains of most residues in the structure can be observed (figs. S6 and S7). The diameter of the ACC1-citrate filaments is about 183 Å (Fig. 1A). Each ACC1 homodimer has a length of 178 Å along the direction of filament extension, contributing to a total length increase of 139 Å for each filament (Fig. 1A, C). There is minimal rotation around the filament extension axis between the two protomers within the same ACC1 homodimer, but a 124° clockwise rotation of the second ACC1 homodimer relative to the first ACC1 homodimer (Fig. 1D).

**Fig. 1.**
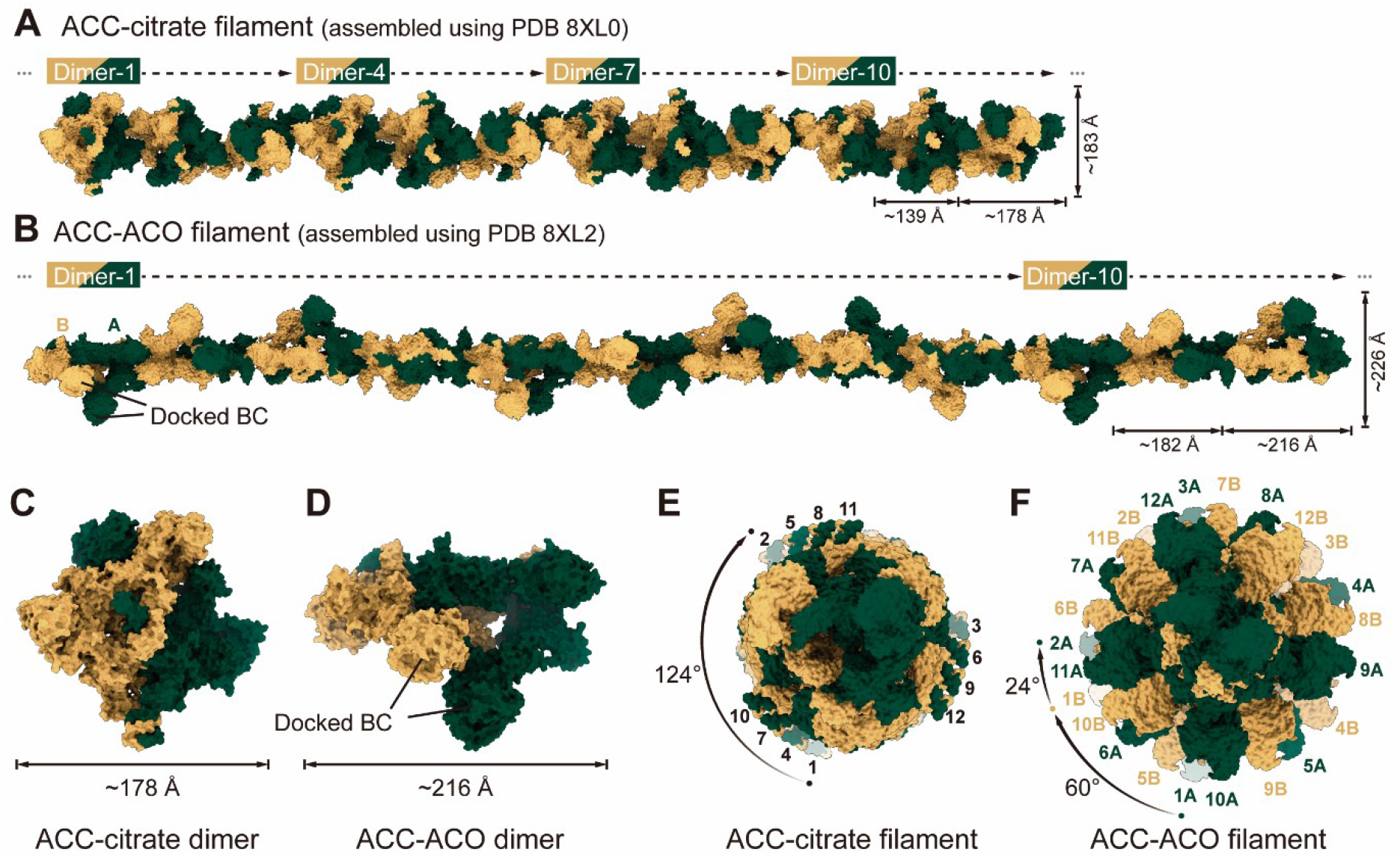
Assembly of human ACC-citrate and ACC-ACO filaments. **(A, B)** Side view of the structural model of the ACC-citrate filaments (**A**) and that of the ACC-ACO filaments (**B**). The cryo-EM structure of the ACC-citrate filament (PDB 8XL0) contains three repeated ACC homodimers, while the cryo-EM structure of the ACC-ACO filament (PDB 8XL2) contains one and a half ACC homodimers. The models were each generated by connecting the cryo-EM structures through the CD nodes (see Fig. 2). The BC domains in the ACC-ACO filament were docked into the structure based on partial density (see fig. S4). Both types of filaments are formed by spirally arranged ACC1 homodimers. Each model shown here contains 12 ACC1 homodimers that are shown in surface representation. The two protomers in each ACC1 homodimer are colored yellow and dark green, respectively. (**C, D**) The structures of the ACC1 homodimer in the ACC-citrate filaments (**C**) and that in the ACC-ACO filaments (**D**). (**E, F**) Top view of the structures of the ACC-citrate filaments (**E**) and the ACC1-ACO filaments (**F**). The diameters of the two types of filaments are approximately 183 Å and 226 Å, respectively. The numbers 1 to 12 refer to the twelve consecutive ACC1 homodimers in each filament, and A and B indicate the two protomers in each ACC1 homodimer.

While the ACC-ACO filaments are also composed of ACC1 homodimers arranged in a stacked manner, their structure exhibits notable differences (Fig. 1B). The ACC-ACO filaments have a diameter of approximately 226 Å. Each ACC1 homodimer extends a length of 216 Å along the direction of filament extension, resulting in a total length increase of 182 Å for each filament (Fig. 1B, E). There is a 60° clockwise rotation of one protomer (protomer B) relative to the other protomer (protomer A) in the same ACC1 homodimer, and a 24° clockwise rotation of protomer A in the second ACC1 homodimer relative to protomer B in the first ACC1 homodimer (Fig. 1F).

### Architectures of the ACC1 homodimers in the ACC filament

The differences between the ACC-citrate filaments and the ACC-ACO filaments are caused by variations in the architecture of the two types of ACC filaments. In both types of filaments, ACC1 homodimers interact with adjacent ACC1 homodimers via their CD domains (Fig. 2A, B). The CD domains from two adjacent ACC1 homodimers form a dimer, called CD node. As previously described(*16*), the CD domain can be divided into four subdomains from N to C termini: CD-N, CD-L, CD-C1, and CD-C2. The interactions between the CD domains in both filaments are facilitated by CD-N of one CD domain interacting with CD-L and CD-C1 of the other CD domain (Fig. 2C, D and fig. S8). However, the CD domains in the ACC-ACO filaments show a conformation different from that in the ACC-citrate filaments. Specifically, when aligning the CD-L and CD-C1 in the two types of filaments, it is observed that the CD-N in the ACC-ACO filaments undergoes a 15° rotation relative to the CD-N in the ACC-citrate filaments.Furthermore, the CD-C2 in the ACC-ACO filaments has a 30° rotation relative to the CD-C2 in the ACC-citrate filaments (Fig. 2E). These findings suggest that the CD domain plays an important role in the dynamic organization of human ACC.

**Fig. 2.**
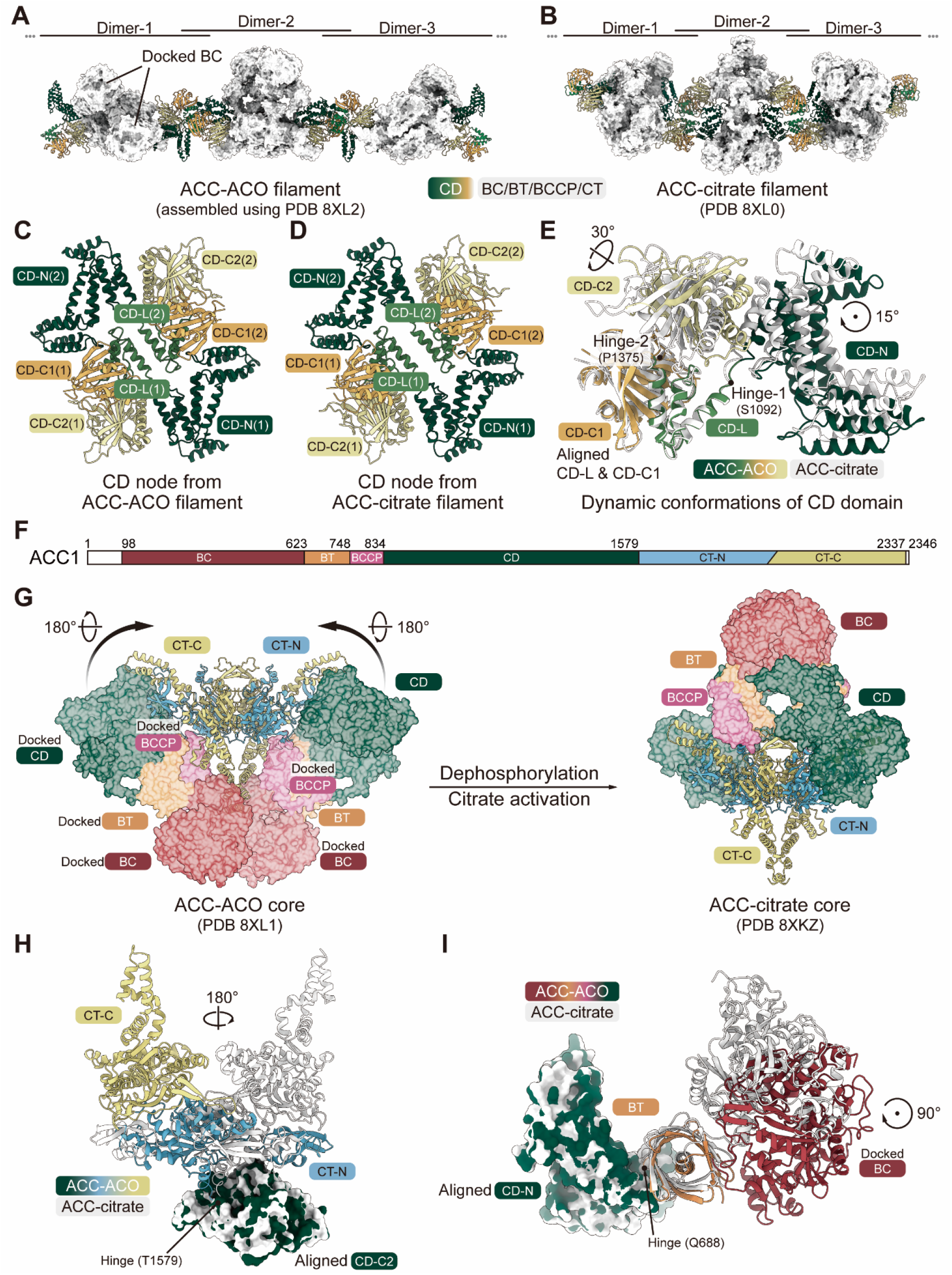
Differences between ACC1 homodimers in the ACC-citrate filaments and the ACC-ACO filaments. (**A, B**) The CD domain mediates the formation of both the ACC-ACO filaments (PDB 8XL2) (**A**) and the ACC-citrate filaments (PDB 8XL0) (**B**). The CD domain is shown in cartoon and colored green to yellow. Other domains are shown in surface and colored gray. (**C, D**) Structures of the CD nodes from the ACC-ACO filaments (**C**) and the ACC-citrate filaments (**D**). The CD-N, CD-C, CD-C1, and CD-C2 subdomains are colored dark green, green, orange, and yellow, respectively. (**E**) Conformational differences between the CD nodes from the ACC-ACO filaments and the ACC-citrate filaments. The CD-L and CD-C1 subdomains are aligned in Chimera X(*32*). The rotation angles of CD-N and CD-C2 in the ACC-ACO filaments relative to those in the ACC-citrate filaments were measured with Hinge-1 (S1092) and Hinge-2 (P1375) as the rotation points. (**F**) Schematic illustration of the domain organization of ACC1. White color indicates the invisible regions. The CT domain consists of CT-N (blue) and CT-C (yellow) subdomains. (**G**) Conformational differences between ACC1 homodimers in the ACC-ACO filaments (ACC-ACO core, PDB 8XL1) and the ACC-citrate filaments (ACC-citrate core, PDB 8XKZ). The CT domains in the two structures adopt a similar conformation, and their structures from the same view are shown as cartoons. The CD and BT in one protomer is invisible in PDB 8XL1; but to show the architecture of the ACC1 homodimer, we docked them in the structure based on the structures of CD and BT in the other protomer. (**H**) The CD-C2 subdomains in the ACC-ACO and the ACC-citrate structures were aligned to show rotation of the CT domains (CT-N and CT-C). (**I**) The CD-N subdomains in the ACC-ACO and the ACC-citrate structures were aligned to show rotation of the BC domains.

In addition to the differences in the conformation of the CD domains, there are also notable differences in the organization of other domains of ACC1 in the two types of ACC filaments (Fig. 2F). The CT domains form dimers at the center of the ACC1 homodimers in both types of ACC filaments. The conformation of the CT dimers in the ACC-ACO filaments is almost identical to that in the ACC-citrate filaments. However, alignment of the CT dimers reveals a 180° rotation of the CD domain in each ACC1 in the ACC-ACO filaments relative to that in the ACC-citrate filaments (Fig. 2G). This rotation results in a distinctly different architecture of the ACC1 homodimer in the ACC-ACO filament compared to that in the ACC-citrate filament (Fig. 2G).

The difference between the relative positions of the CD and BC domains in the two types of filaments was also illustrated by aligning the CD domains, which shows a 180° difference between the CT domains (Fig. 2H). The alignment of the CD domains also demonstrates a 90° rotation of the BC domains in ACC-ACO filaments in comparison to those in the ACC-citrate filaments (Fig. 2I).

### An inactive conformation of ACC1 in the ACC-ACO filaments

As reported previously, the ACC1 homodimers in the ACC-citrate filaments are in an active state (Fig. 3A)(*16*). The two BC domains in each ACC1 homodimer dimerize to adopt an active conformation. The active site of the BC domain is approximately 80 Å away from the active site of the CT domain. The BCCP domain, in which K786 is covalently linked with biotin, is close to the CT domain. A rotation of about 105° of the BCCP domain can switch the biotin from the CT domain to the active site in the BC domain. The switch of the BCCP domain between the BC and CT domains facilitates the two-step carboxylation reaction(*8, 16*).

**Fig. 3.**
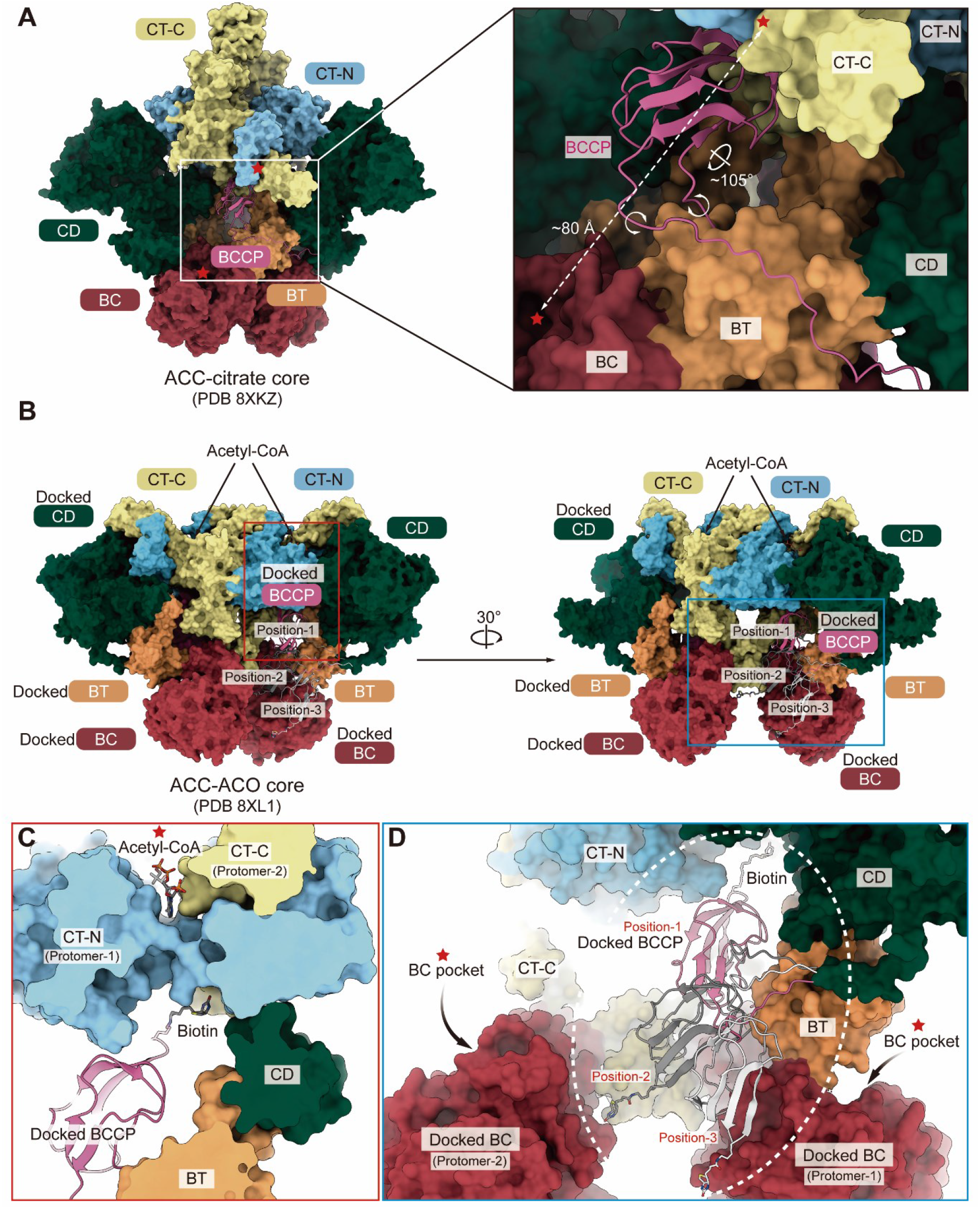
ACC1 in the ACC-ACO filaments adopts an inactive conformation. (**A**) The structure of the ACC1 homodimer in the ACC-citrate filaments (ACC-citrate core, PDB 8XKZ). The BCCP domain, which is shown in cartoon representation, binds to the CT dimer interface. During catalysis, the BCCP domain swings between the CT and BC domains. The dashed arrow indicates the distance between BC and CT reaction centers. A 105° rotation of the BCCP domain around the connecting linkers can switch the BCCP domain from the CT domain to the BC domain. (**B**) The structure of the ACC1 homodimer in the ACC-ACO filaments (ACC-ACO core, PDB 8XL1). The structural model shown in the figure was generated as described in Fig. 2G. Three possible locations of the BCCP domain were generated according to the positions of the BT and CD domains. (**C**) The relative position of the biotin and the acetyl-CoA in the ACC-ACO filaments when the covalently linked biotin in the BCCP domain is placed at the position closest to the acetyl-CoA binding site. The biotin and the acetyl-CoA are separated by the CT domain. (**D**) The relative position of the biotin and the active sites of the BC domains in the ACC-ACO filaments. The white dotted circle indicates the possible range of movement of the biotin covalently linked to the BCCP domain.

In contrast, in the ACC-ACO filaments, the two BC domains in each ACC1 homodimer are in a monomeric state and separate from each other (Fig. 3B), indicating that ACC1 in the ACC-ACO filaments is in an inactive state. The BCCP domain was not resolved in the cryo-EM map, probably due to its flexibility (fig. S4). However, the BT and CD domains, which are attached to the N and C termini of the BCCP domain, were clearly resolved. To examine whether the BCCP domain is able to access the acetyl-CoA binding site, we docked the biotinylated BCCP domain into the structure based on the positions of the BT and CD domains (Fig. 3B and fig. S4). The BCCP domain is blocked by the CT-N domain from accessing the acetyl-CoA binding site (Fig. 3C). Moreover, even when considering the maximum range of possible movement of the BCCP domain, the BCCP domain remains unable to access either of the BC domains in the ACC1 homodimer (Fig. 3D). These findings further confirm that ACC1 in the ACC-ACO filaments is in an inactive state.

### Binding of the covalently linked biotin to an exo-site in the ACC-citrate filaments

In our structure of the ACC-citrate filaments, there is a biotin covalently linked to each BCCP domain. The biotin binds to a pocket formed by a CT-N and two CT-C subdomains (Fig. 4A). The two amino groups of the biotin donate two hydrogen bonds to D2000 from one CT-C and S2050 from the other CT-C. The ureido ring carbonyl oxygen and the carboxyl group of the biotin accept hydrogen bonds from the main chain amide of D2055 and the side chain of R2047, respectively (Fig. 4A).

**Fig. 4.**
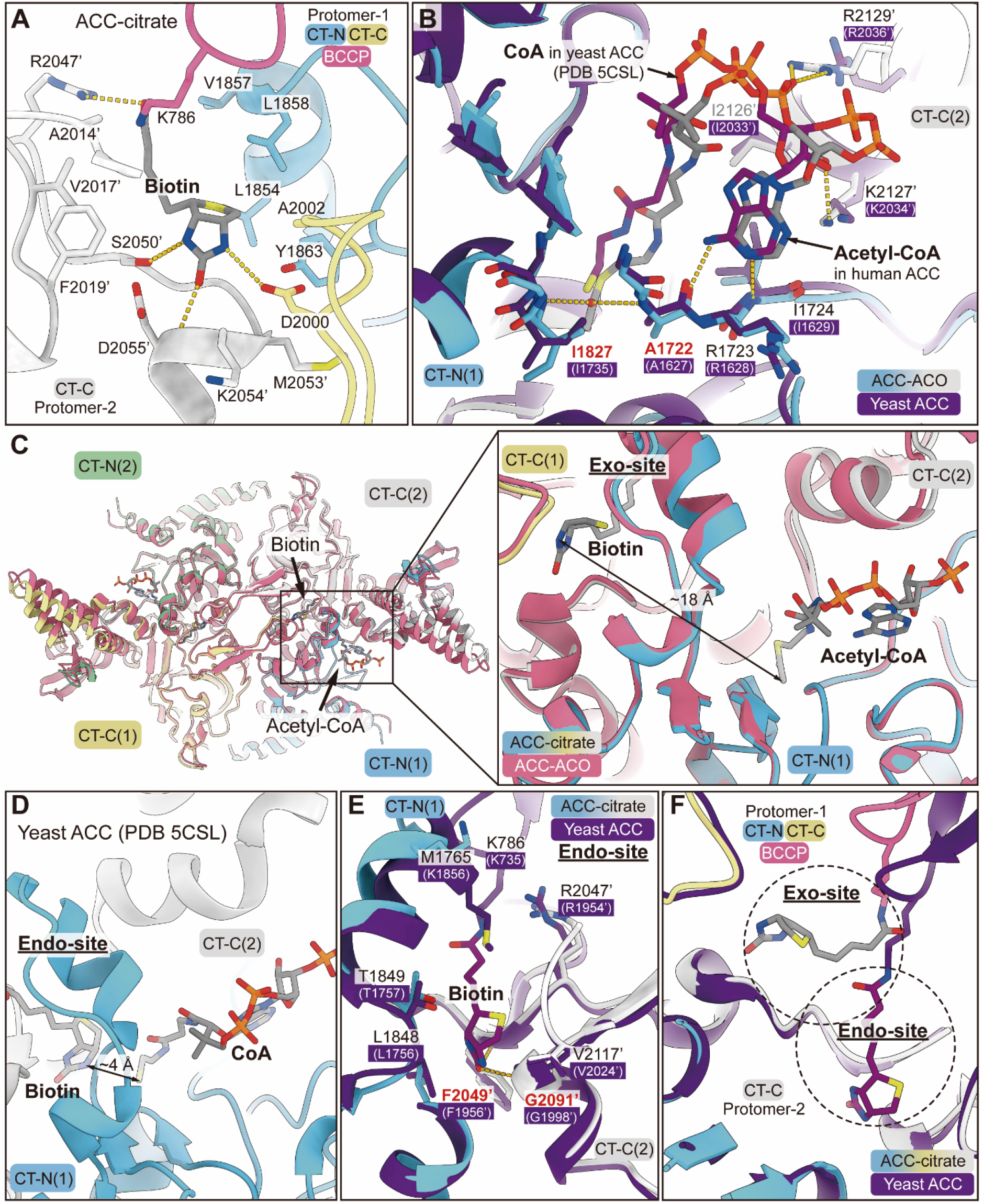
Biotin binds at an exo-site in the CT domain of human ACC1. (**A**) Interactions between the covalently linked biotin and ACC1 in the ACC-ACO filaments. The CT-N and CT-C subdomains, and the BCCP domain are colored blue, yellow, and pink, respectively. The residue numbers with a primer (‘) indicate residues from the other protomer in a ACC1 homodimer. (**B**) Alignment of the CT domain in the ACC-ACO structure with that in the yeast ACC structure (PDB code: 6G2D) shows that acetyl-CoA binds to similar sites in the two structures. (**C**) Alignment of the CT domain in the ACC-ACO structure with that in the ACC-citrate structure. The carboxylation site of the covalently linked biotin in the ACC-citrate structure is about 18 Å far away from the carboxylation site of acetyl-CoA. The biotin binding site observed in the ACC-citrate structure is thus referred to as the “exo-site”. (**D**) In the yeast ACC structure (PDB code: 6G2D), the distance between the carboxylation site of the covalently linked biotin and that of acetyl-CoA is about 4 Å. The biotin binding site observed in the yeast ACC structure is referred to as the “endo-site”. (**E, F**) Alignment of the CT domain in the ACC-citrate structure with that in the yeast ACC structure (PDB code: 6G2D). The residues form the endo-site for biotin binding in yeast ACC, and the corresponding residues in human ACC1 are shown as sticks (**E**). The exo-site and the endo-site are located in separate positions (**F**).

In the structure of the ACC-ACO filaments, no biotin is observed, but an acetyl-CoA is observed to bind to a pocket formed by the CT-N of one protomer and CT-C of the other protomer in each ACC1 homodimer (Fig. 4B). The binding mode of acetyl-CoA in the ACC-ACO filaments is similar to that in the yeast ACC homodimer (PDB code: 5CSL)(*8*).

Alignment of the CT dimer in the ACC-citrate filaments with that in the ACC-ACO filaments indicates that there is a distance of 18 Å between the N1 of biotin, where the carboxylation reaction occurs, and the Cα of acetyl-CoA, where the carboxyl group is transferred to (Fig. 4C). This distance exceeds the range required for the transfer of a carboxyl group from biotin to acetyl-CoA. In comparison, in the yeast ACC homodimer, the distance between the carboxylation sites of biotin and acetyl-CoA is approximately 4 Å (Fig. 4D). In the yeast ACC, the heterocyclic ring of biotin binds to a pocket formed by residues L1756, T1757, R1954 to F1956, G1998, and V2024, which corresponds to residues L1848, T1849, R2047 to F2049, G2091, and V2117 in human ACC1 (Fig. 4E)(*8*).

When aligning the CT dimer in the ACC-citrate homodimer with that in the yeast ACC homodimer, it becomes evident that biotin occupies two distinct sites: the site near the acetyl-CoA binding site is referred to as the endo-site, while the site far from the acetyl-CoA binding site is called the exo-site (Fig. 4F). Binding of the covalently linked biotin to the endo-site facilitates the transfer of a carboxyl group from biotin to acetyl-CoA. In contrast, biotin binding to the exo-site cannot catalyze the carboxylation reaction, suggesting it may represent a resting state of the covalently linked biotin.

## Discussion

Two types of human ACC filaments were reported before our study(*16*): one is the ACC-citrate filaments, and the other is the ACC-BRCT filaments, which represent an active state and an inhibited state of ACC1, respectively. The cryo-EM structures solved in our study provide more details on the ACC-citrate filaments and, more importantly, reveal a new type of ACC filaments: the ACC-ACO filaments that represent an inactive state of ACC1.

Our findings highlight the dynamic organization of human ACC1. An important question is: how does ACC1 transform between the different filament states? Based on the cryo-EM structures of the ACC-citrate filaments and the ACC-BRCT filaments, Moritz Hunkeler *et al*. proposed that one type of ACC1 filaments needs to be dissociated into ACC1 homodimers first. Then, upon phosphorylation or dephosphorylation, and association or dissociation with ACC1 regulators (citrate, BRCT, etc.), the ACC1 homodimers can be stabilized into a new conformation, thus facilitating the transformation into a second type of ACC1 filaments(*16*). This model can also be used to describe the transition between the ACC-ACO filaments and the ACC-citrate filaments. However, it cannot explain why ACC1 forms such inactive filaments but does not simply remain as inactive homodimers. One possibility is that the ACC-ACO filaments can be directly converted into the ACC-citrate filaments, allowing ACC1 to be activated in a highly efficient manner. Both the ACC-ACO and ACC-citrate filaments are comprised of ACC1 homodimers, which are arranged by stacking one another through their CD domains. The transition from the inactive ACC-ACO filaments to the active ACC-citrate filaments may involve a 180° rotation of the CD domain relative to the CT domain, followed by a 90° rotation of the BC domain (Fig. 2). The plasticity of the CD nodes, as illustrated by the alignment of the structure of the CD domains in the ACC-ACO filaments with that in the ACC-citrate filaments (Fig. 2), may facilitate the structural transition while maintaining the overall filament architecture.

The structure of the ACC-citrate filaments has a resolution of 2.55 Å. However, the binding of citrate in the structure still cannot be observed. In the crystal structure of full-length yeast ACC, 0.2 M sodium citrate was included in the crystallization buffer, but no citrate binding was observed in the structure(*8*). As suggested by Jia Wei and Liang Tong, citrate may play a nonspecific role in activating both yeast and human ACCs(*11*). But the molecular mechanism is still unclear.

Another interesting finding in our study is that each covalently linked biotin in the ACC-citrate filaments binds to an exo-site that is far away from the corresponding acetyl-CoA binding site (Fig. 4). An exo-site (or “exo pocket”) for biotin was first identified in *Staphylococcus aureus* PYC(*20*). Biotin was also found to bind at the pocket far away from the catalytic sites in the CT domains of ACC, PCC and MCC, indicating the conservation of the exo-site in BDC family(*8,21,22*). According to the well-accepted two-step reaction mechanism for the carboxylation of acetyl-CoA, the covalently linked biotin is first carboxylated in the BC domain and then translocated to the CT domain to transfer the carboxyl group to acetyl-CoA(*1,6,23*). Biotin binding to the exo-site cannot transfer its carboxyl group to acetyl-CoA, thus it is in a catalytically incompetent state. One possible explanation is that in the absence of acetyl-CoA, the biotin molecule occupies the exo-site as a resting state. However, in the presence of acetyl-CoA, there is coordination between acetyl-CoA binding and biotin binding. As a result, the biotin occupies the endo-site to facilitate the carboxylation reaction.

## Materials and Methods

### Cell culture

Expi 293F cells (Thermo Scientific) were cultured in SMM 293T-II medium (Sino Biological) at 37 °C with 5% CO_2_ in a Multitron-Pro incubator (Infors). After the density reached 2.0×10^6^ per mL, the cells were cultured for an additional 48 h, then harvested by centrifuging at 4,000 ×*g* (Fiberlite F9-6×1000 LEX rotor, Thermo Scientific) for 10 min and resuspended with HBS buffer (20 mM HEPES, pH 7.4, 150 mM NaCl). The cells were snap frozen and stored at −80 °C.

### Endogenous purification of biotin dependent carboxylases

The frozen cells were thawed and disrupted using a high-pressure cell crusher and centrifuged at 23,376 ×*g* (JA-14.50 rotor, Beckman) for 1 h at 4 °C. The supernatant was incubated with Strep-Tactin^®^XT resin (IBA Lifesciences) for 1 h at 4 °C and then washed with HBS buffer. The proteins were eluted by HBS buffer supplemented with 50 mM *D*-biotin (J&K Scientific) and further purified by size-exclusion chromatography (Superose 6 Increase 10/300 GL column, Cytiva) with HBS as the running buffer. For dephosphorylation, the fractions eluted within 12– 13.5 mL were pooled and concentrated to 4 mg/mL, then incubated with a 1:5 molar ratio of purified PP1α at 4 °C overnight with the presence of 2 mM MnCl_2_. The sample was then dialyzed against HBS buffer supplemented with 10 mM sodium citrate (Sigma-Aldrich) at 4 °C for 6 h, yielding the ACC-citrate sample with a final concentration of 0.42 mg/mL. For the substrates-bound condition, another batch of proteins were purified accordingly, and fractions eluted within 12.5–16.5 mL were pooled and concentrated to 2.65 mg/mL. The sample was pre-incubated with 25 mM NaHCO_3_, 10 mM MgCl_2_ and 10 mM acetyl-CoA for 1 h on ice, and then further incubated with 10 mM of ATP for an additional 1 h at room temperature, yielding the ACC-ACO sample. The proteins were analyzed by mass spectrometry and applied for cryo-EM grids preparation the same day purified.

### Cryo-EM data acquisition and pre-processing

Lacey grids (Lacey carbon filmed 400 mesh cooper grids, Ted Pella) or Holey grids (Holey carbon filmed 300 mesh gold grids, R1.2/1.3, Quantifoil) were glow discharged with a medium RF power for 30 s using a plasma cleaner (PDC-32G-2, Harrick Plasma) prior use. The ACC-citrate sample was incubated with 0.01% (*w/v*) DDM (Anatrace) for 1 h at room temperature. Subsequently, 4 μL of the sample was loaded to a Lacey grid and then flash frozen in liquid ethane immediately using Vitrobot (Mark IV, Thermo Fisher Scientific). For the ACC-ACO sample, the pre-treated proteins were directly frozen on a Holey grid. The cryo-EM grids were stored in liquid nitrogen until data acquisition.

The cryo-EM grids were transferred to a Titan Krios TEM (FEI) operating at 300 kV equipped with a K3 Summit direct electron detector (Gatan) and a GIF Quantum energy filter (Gatan). Zero-loss movie stacks were automatically collected using EPU software (FEI) with a slit width of 20 eV on the energy filter and a defocus range from −1.2 to −2.2 μm in super-resolution mode at an 81,000× nominal magnification. Each movie stack, which contained 32 frames, was exposed for 2.56 s with a total electron dose of ∼50 e^-^/Å^2^. The stacks were motion corrected using MotionCor2 with a binning factor of 2, resulting in the pixel size of 1.0773 Å(*24*). Dose weighting was performed concurrently(*25*). The movie stacks were imported to cryoSPARC (v3.3.2, Structura Biotechnology) for CTF estimation and downstream processing(*26*).

### Cryo-EM image processing

For the ACC-citrate sample, 4,669 motion corrected micrographs were imported to cryoSPARC. After patch CTF estimation, a total of 586,745 particles were automatically picked according to the templates generated from an initial 2D classification of manually picked particles. The particles were extracted with a box size of 488 pixels. After a round of 2D classification with 20 online-EM iterations, the dataset was sorted into 400 classes, 37 of which with identifiable features were selected, yielding 144,582 “good” particles. Three maps were randomly generated using the selected particles, and three using the excluded “bad” particles. A heterogeneous refinement job was carried out using these initial maps, yielding more detailed maps that shown clearer secondary structure features. The non-uniform refinement job was subjected using the selected 3D volume and corresponding particles, yielding a 4.14 Å map of ACC-citrate filaments. To improve the resolution of the core region of ACC-citrate filaments, a template-based picking was performed using 26 equally spaced templates generated with the 4.14 Å map. 2D classification was performed with 549,746 re-extracted (box size = 320 pixels) particles and the “good” classes including 354,467 particles were selected, which were then subjected to a simultaneous ab-initio reconstruction of multiple volumes, yielding a map with clearer features composed by 81.2% of the particles used. Heterogenous refinement was performed subsequently after flipping the chirality of the initial map by the “*relion_image_handler*” implement(*27*). The improved map was then polished, generating the final map of the core region of ACC-citrate filament (ACC-citrate core) at 2.55 Å resolution.

For the ACC-ACO sample, 13,632 micrographs were collected and imported to cryoSPARC after preprocessing. Particles were automatically picked and extracted (box size = 488 pixels) as described above. After an initial round of 2D classification, the selected particles were further classified and separated into two clusters composed of filamentous (cluster-A, 689,638 particles) and granular (cluster-B, 461,070 particles) particles, respectively. The filamentous particles within cluster-A were subjected to ab-initio reconstruction and heterogeneous refinement jobs, gathering 36% of the particles and producing the refined map of ACC filaments. These particles were re-extracted with different box sizes (488 or 320 pixels) and subjected to non-uniform refinement jobs, respectively, yielding the maps of ACC-ACO filaments at 2.73 Å and its core region (ACC-ACO core) at 2.57 Å.

The resolutions mentioned above were determined according to the gold-standard Fourier shell correlation 0.143 criterion and with a high-resolution noise substitution method(*28*). All the reconstructed maps were subjected to further model building.

### Model building and structure refinement

The reported structure (PDB code: 6G2D) was used as a template for the model building of ACC filaments(*16*). The reference model was automatically placed using Rosetta and manually adjusted using Coot based on the cryo-EM maps, resulting the modified model(*29, 30*). Each residue was manually checked with the chemical properties taken into consideration during model building. Several segments, whose corresponding densities were invisible, were not modeled.

Real space refinement was performed using Phenix with secondary structure and geometry restraints to prevent overfitting(*31*). For the cross-validation of the structure, the model was refined against one of the two independent half maps from the gold-standard 3D refinement approach. Then, the refined model was tested against the other half map. Statistics associated with data collection, 3D reconstruction and model building were summarized in Table S2.

## Acknowledgments

We thank Drs. Xiaofeng Zhang and Xiechao Zhan for valuable discussion and suggestions on structure determination. We thank the Cryo-EM Facility, the High-Performance Computing Center, and the Mass Spectrometry & Metabolomics Core Facility of Westlake University for technical support.

## Funding

“Pioneer” and “Leading Goose” R&D Program of Zhejiang 2024SSYS0036 (Q.H.)

Westlake Laboratory of Life Sciences and Biomedicine (Q.H.)

Westlake Education Foundation (Q.H.)

## Author contributions

Conceptualization: Q.H., F.Z

Methodology: Q.H., F.Z.

Investigation: F.Z., Y.Zhang, Y.Zhu

Visualization: F.Z., Y.Zhang

Funding acquisition: Q.H.

Project administration: Q.H., F.Z.

Supervision: Q.H., Y.S.

Writing – original draft: Q.H., F.Z.

Writing – review & editing: Q.H., Y.S., Q.Z.

## Data and materials availability

All data needed to evaluate the conclusions in the paper are present in the paper and the Supplementary Materials. The cryo-EM structures have been deposited in the Protein Data Bank with the accession codes 8XKZ (ACC-citrate core), 8XL0 (ACC-citrate filament), 8XL1 (ACC-ACO core) and 8XL2 (ACC-ACO filament). The cryo-EM maps have been deposited in the Electron Microscopy Data Bank with the accession codes EMD-38432 (ACC-citrate core), EMD-38433 (ACC-citrate filament), EMD-38434 (ACC-ACO core) and EMD-38435 (ACC-ACO filament). The crystal structure of the yeast ACC (PDB code: 5CSL), and the cryo-EM structure of the human ACC-citrate filaments (PDB code: 6G2D) were used for structural analysis in this study. Human ACC1 and ACC2 sequences used in this study are available from the Universal Protein Knowledgebase (UniProtKB) with the accession codes Q13085 and O00763, respectively.

## Supplementary materials

### The PDF file includes

Figs. S1 to S8 Tables S2

### Other Supplementary Material for this manuscript includes the following

Tables S1

## References

1. L. Tong, Structure and function of biotin-dependent carboxylases. Cell Mol Life Sci 70, 863–891 (2013).

2. S. J. Wakil, J. K. Stoops, V. C. Joshi, Fatty acid synthesis and its regulation. Annu Rev Biochem 52, 537–579 (1983).

3. J. E. Cronan, Jr., G. L. Waldrop, Multi-subunit acetyl-CoA carboxylases. Prog Lipid Res 41, 407–435 (2002).

4. A. Chapman-Smith, J. E. Cronan, Jr., Molecular biology of biotin attachment to proteins. J Nutr 129, 477S–484S (1999).

5. J. R. Knowles, The mechanism of biotin-dependent enzymes. Annu Rev Biochem 58, 195–221 (1989).

6. L. Tong, in Adv Protein Chem Struct Biol. (2017), vol. 109, pp. 161–194.

7. C.-Y. Chou, L. P. C. Yu, L. Tong, Crystal structure of biotin carboxylase in complex with substrates and implications for its catalytic mechanism. J Biol Chem 284, 11690–11697 (2009).

8. J. Wei, L. Tong, Crystal structure of the 500-kDa yeast acetyl-CoA carboxylase holoenzyme dimer. Nature 526, 723–727 (2015).

9. U. Hoja, S. Marthol, J. Hofmann, S. Stegner, R. Schulz, S. Meier, E. Greiner, E. Schweizer, HFA1 encoding an organelle-specific acetyl-CoA carboxylase controls mitochondrial fatty acid synthesis in Saccharomyces cerevisiae. J Biol Chem 279, 21779–21786 (2004).

10. K. W. Kim, H. Yamane, J. Zondlo, J. Busby, M. Wang, Expression, purification, and characterization of human acetyl-CoA carboxylase 2. Protein Expr Purif 53, 16–23 (2007).

11. J. Wei, L. Tong, How Does Polymerization Regulate Human Acetyl-CoA Carboxylase 1? Biochemistry 57, 5495–5496 (2018).

12. J. Wei, Y. Zhang, T.-Y. Yu, K. Sadre Bazzaz, M. J. Rudolph, G. A. Amodeo, L. S. Symington, T. Walz, L. Tong, A unified molecular mechanism for the regulation of acetyl-CoA carboxylase by phosphorylation. Cell Discov 2, 16044 (2016).

13. C. W. Kim, Y. A. Moon, S. W. Park, D. Cheng, H. J. Kwon, J. D. Horton, Induced polymerization of mammalian acetyl-CoA carboxylase by MIG12 provides a tertiary level of regulation of fatty acid synthesis. Proc Natl Acad Sci U S A 107, 9626–9631 (2010).

14. R. W. Brownsey, A. N. Boone, J. E. Elliott, J. E. Kulpa, W. M. Lee, Regulation of acetyl-CoA carboxylase. Biochem Soc Trans 34, 223–227 (2006).

15. G. A. Locke, D. Cheng, M. R. Witmer, J. K. Tamura, T. Haque, R. F. Carney, A. R. Rendina, J. Marcinkeviciene, Differential activation of recombinant human acetyl-CoA carboxylases 1 and 2 by citrate. Arch Biochem Biophys 475, 72–79 (2008).

16. M. Hunkeler, A. Hagmann, E. Stuttfeld, M. Chami, Y. Guri, H. Stahlberg, T. Maier, Structural basis for regulation of human acetyl-CoA carboxylase. Nature 558, 470–474 (2018).

17. R. W. Brownsey, N. J. Edgell, T. J. Hopkirk, R. M. Denton, Studies on insulin-stimulated phosphorylation of acetyl-CoA carboxylase, ATP citrate lyase and other proteins in rat epididymal adipose tissue. Evidence for activation of a cyclic AMP-independent protein kinase. Biochemical Journal 218, 733–743 (1984).

18. J. Ha, S. Daniel, S. S. Broyles, K. H. Kim, Critical phosphorylation sites for acetyl-CoA carboxylase activity. J Biol Chem 269, 22162–22168 (1994).

19. C. Magnard, R. Bachelier, A. Vincent, M. Jaquinod, S. Kieffer, G. M. Lenoir, N. D. Venezia, BRCA1 interacts with acetyl-CoA carboxylase through its tandem of BRCT domains. Oncogene 21, 6729–6739 (2002).

20. S. Xiang, L. Tong, Crystal structures of human and Staphylococcus aureus pyruvate carboxylase and molecular insights into the carboxyltransfer reaction. Nat Struct Mol Biol 15, 295–302 (2008).

21. J. K. J. Lee, Y. T. Liu, J. J. Hu, I. Aphasizheva, R. Aphasizhev, Z. H. Zhou, CryoEM reveals oligomeric isomers of a multienzyme complex and assembly mechanics. J Struct Biol X 7, 100088 (2023).

22. J. J. Hu, J. K. J. Lee, Y. T. Liu, C. Yu, L. Huang, I. Aphasizheva, R. Aphasizhev, Z. H. Zhou, Discovery, structure, and function of filamentous 3-methylcrotonyl-CoA carboxylase. Structure 31, 100–110 e104 (2023).

23. J. Lombard, D. Moreira, Early evolution of the biotin-dependent carboxylase family. BMC Evol Biol 11, 232 (2011).

24. S. Q. Zheng, E. Palovcak, J. P. Armache, K. A. Verba, Y. Cheng, D. A. Agard, MotionCor2: anisotropic correction of beam-induced motion for improved cryo-electron microscopy. Nat Methods 14, 331–332 (2017).

25. T. Grant, N. Grigorieff, Measuring the optimal exposure for single particle cryo-EM using a 2.6 A reconstruction of rotavirus VP6. Elife 4, e06980 (2015).

26. A. Punjani, J. L. Rubinstein, D. J. Fleet, M. A. Brubaker, cryoSPARC: algorithms for rapid unsupervised cryo-EM structure determination. Nat Methods 14, 290–296 (2017).

27. S. H. Scheres, A Bayesian view on cryo-EM structure determination. J Mol Biol 415, 406–418 (2012).

28. S. Chen, G. McMullan, A. R. Faruqi, G. N. Murshudov, J. M. Short, S. H. Scheres, R. Henderson, High-resolution noise substitution to measure overfitting and validate resolution in 3D structure determination by single particle electron cryomicroscopy. Ultramicroscopy 135, 24–35 (2013).

29. R. F. Alford, A. Leaver-Fay, J. R. Jeliazkov, M. J. O’Meara, F. P. DiMaio, H. Park, M. V. Shapovalov, P. D. Renfrew, V. K. Mulligan, K. Kappel, J. W. Labonte, M. S. Pacella, R. Bonneau, P. Bradley, R. L. Dunbrack, Jr., R. Das, D. Baker, B. Kuhlman, T. Kortemme, J. J. Gray, The Rosetta All-Atom Energy Function for Macromolecular Modeling and Design. J Chem Theory Comput 13, 3031–3048 (2017).

30. P. Emsley, B. Lohkamp, W. G. Scott, K. Cowtan, Features and development of Coot. Acta Crystallogr D Biol Crystallogr 66, 486–501 (2010).

31. P. D. Adams, P. V. Afonine, G. Bunkoczi, V. B. Chen, I. W. Davis, N. Echols, J. J. Headd, L. W. Hung, G. J. Kapral, R. W. Grosse-Kunstleve, A. J. McCoy, N. W. Moriarty, R. Oeffner, R. J. Read, D. C. Richardson, J. S. Richardson, T. C. Terwilliger, P. H. Zwart, PHENIX: a comprehensive Python-based system for macromolecular structure solution. Acta Crystallogr D Biol Crystallogr 66, 213–221 (2010).

32. E. C. Meng, T. D. Goddard, E. F. Pettersen, G. S. Couch, Z. J. Pearson, J. H. Morris, T. E. Ferrin, UCSF ChimeraX: Tools for structure building and analysis. Protein Sci 32, e4792 (2023).

