## Supplementary Materials for "Filament structures unveil the dynamic organization of human acetyl-CoA carboxylase"

### Supplementary Figures

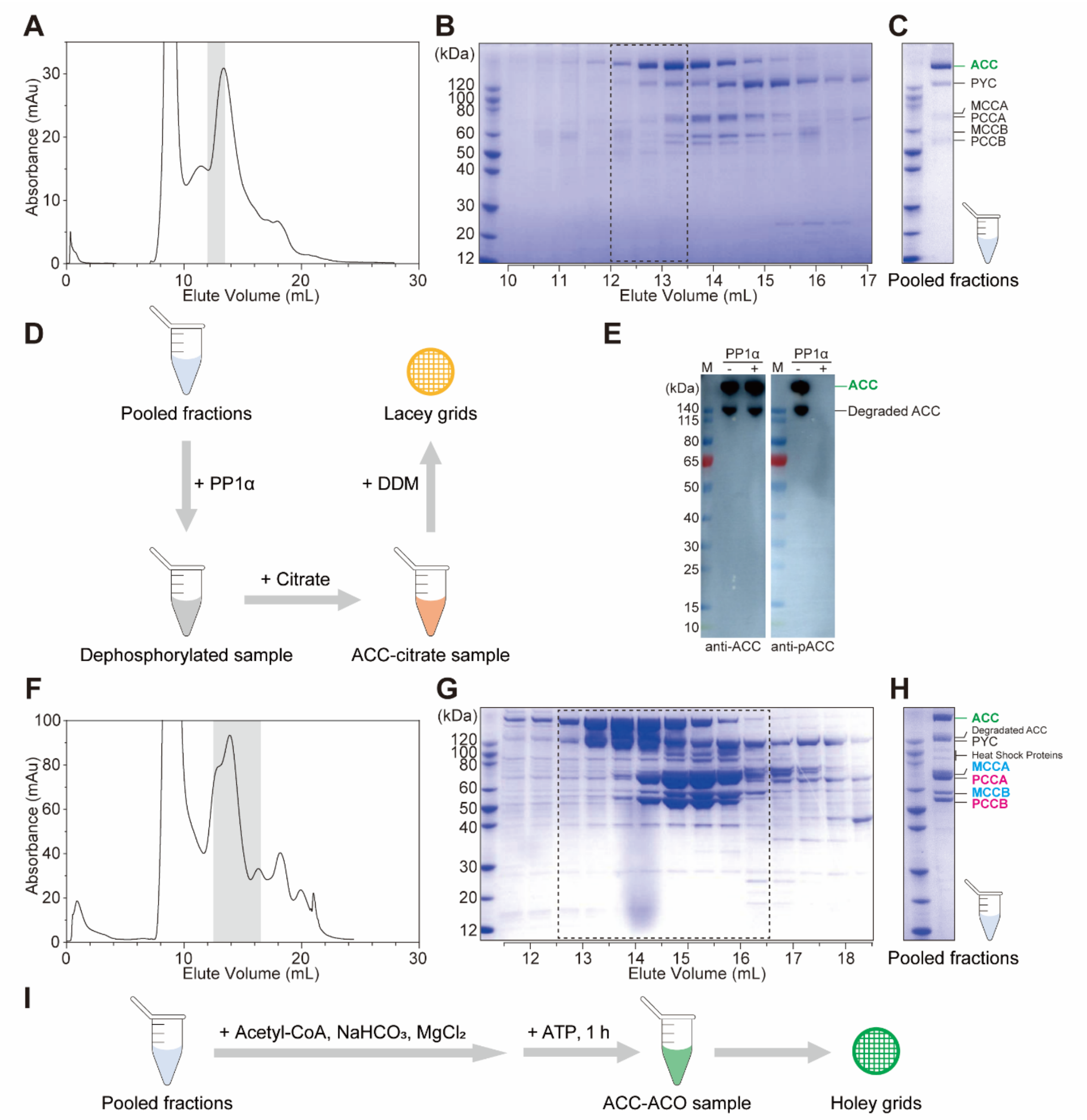


**Fig. S1. Protein purification and cryo-EM sample preparation.** (**A**–**D**) Preparation of the ACC-citrate sample. The proteins eluted from Strep-Tactin®XT resin was further purified by size-exclusion chromatography (**A**). The fractions were analyzed by SDS-PAGE and visualized by Coomassie blue staining (**B**). The peak fractions in which ACC was the major component were combined and analyzed by mass spectrometry (**C**). To prepare the cryo-EM sample, the purified ACC1 was dephosphorylated by PP1α, treated with citrate, and after incubation with DDM, the sample was loaded onto a Lacey Carbon filmed 400-mesh copper grid (Ted Pella) and frozen in liquid ethane (**D**). (**E**) After treatment with PP1α, the sample was analyzed by Western blot. Antibodies against ACC (anti-ACC, CST#3676) and phosphorylated ACC (anti-pACC, CST#11818) were used to detect the total ACC and S80 phosphorylated ACC, respectively. (**F**–

**I**) Preparation of the ACC-ACO sample. The proteins eluted from Strep-Tactin®XT resin was further purified by size-exclusion chromatography (**F**). The fractions were analyzed by SDS-PAGE and visualized by Coomassie blue staining (**G**). The peak fractions were combined and analyzed by mass spectrometry (**H**). To prepare the cryo-EM sample, the purified proteins were concentrated, mixed with acetyl-CoA, $NaHCO_3$, and $MgCl_2$, and then incubated with ATP for 1 hour before frozen on a Holey Carbon filmed 300-mesh gold grid (Quantifoil) (**I**).

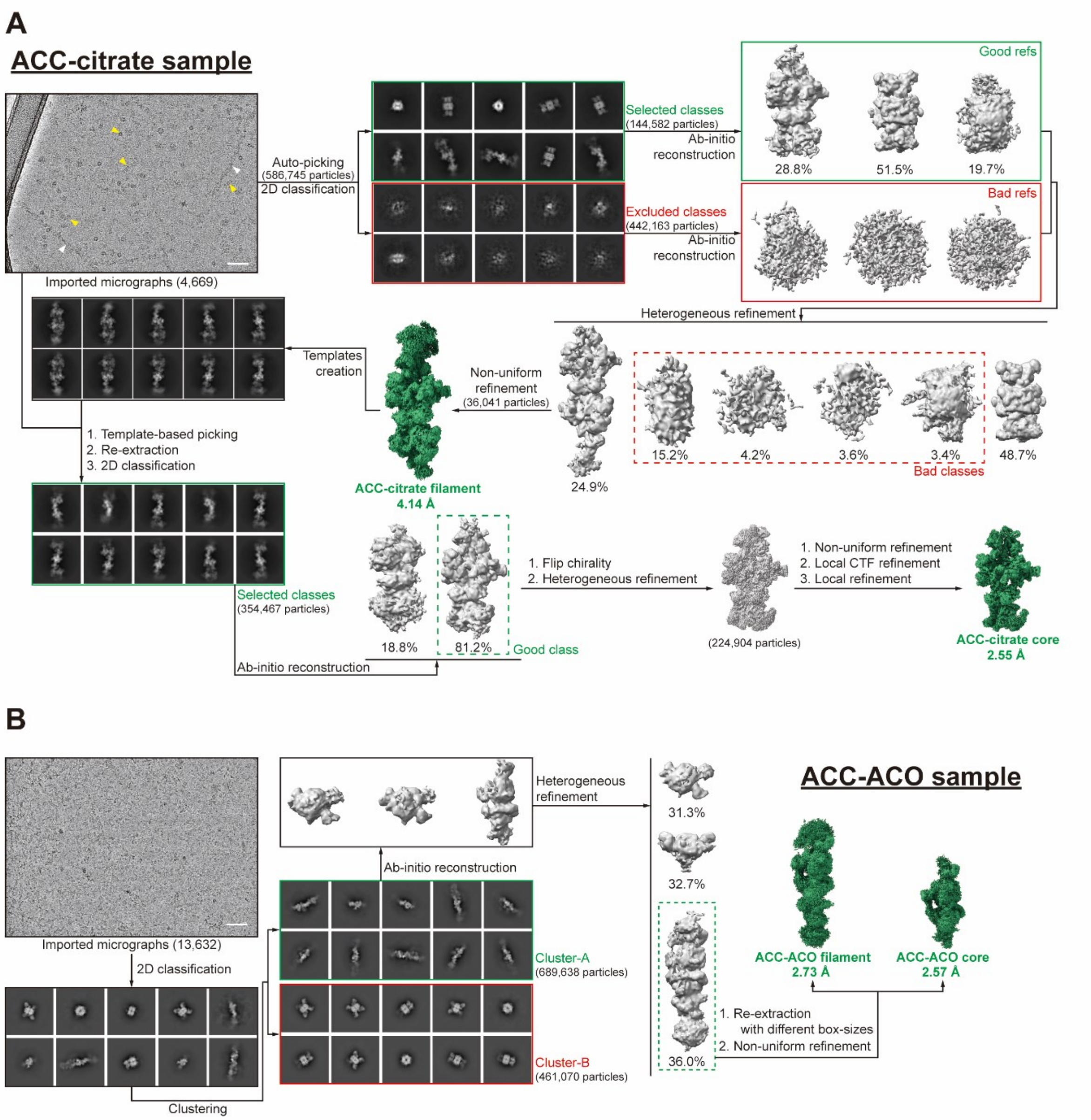


**Fig. S2. Flowchart for cryo-EM data processing of human ACC filaments.** (**A**) Cryo-EM data processing of the ACC-citrate filaments. A total of 586,745 particles were picked from 4,669 micrographs. After multiple rounds of classification, several EM density maps were generated. The particles used to generate the map of the ACC-citrate filaments account for 24.9% of all particles. The density map of the ACC-citrate filaments has a resolution of 4.14 Å. The data was further processed, focusing on the core region of the filaments (named ACC-citrate core), to achieve a density map with a resolution of 2.55 Å. The white triangles and the yellow triangles in the microscopic photo indicate ACC-citrate filaments and particles of other biotin-dependent carboxylases, respectively. (**B**) Cryo-EM data processing of the ACC-ACO filaments. The particles derived from 13,632 micrographs were selected to reconstruct the cryo-EM maps. The particles used to generate the map of the ACC-ACO filaments account for 36% of tubular particles in cluster-A. The final reconstructed map of the ACC-ACO filaments has a resolution of 2.73 Å, and its core region (ACC-ACO core) has a resolution of 2.57 Å. The scale bar in each micrograph is 50 nm.

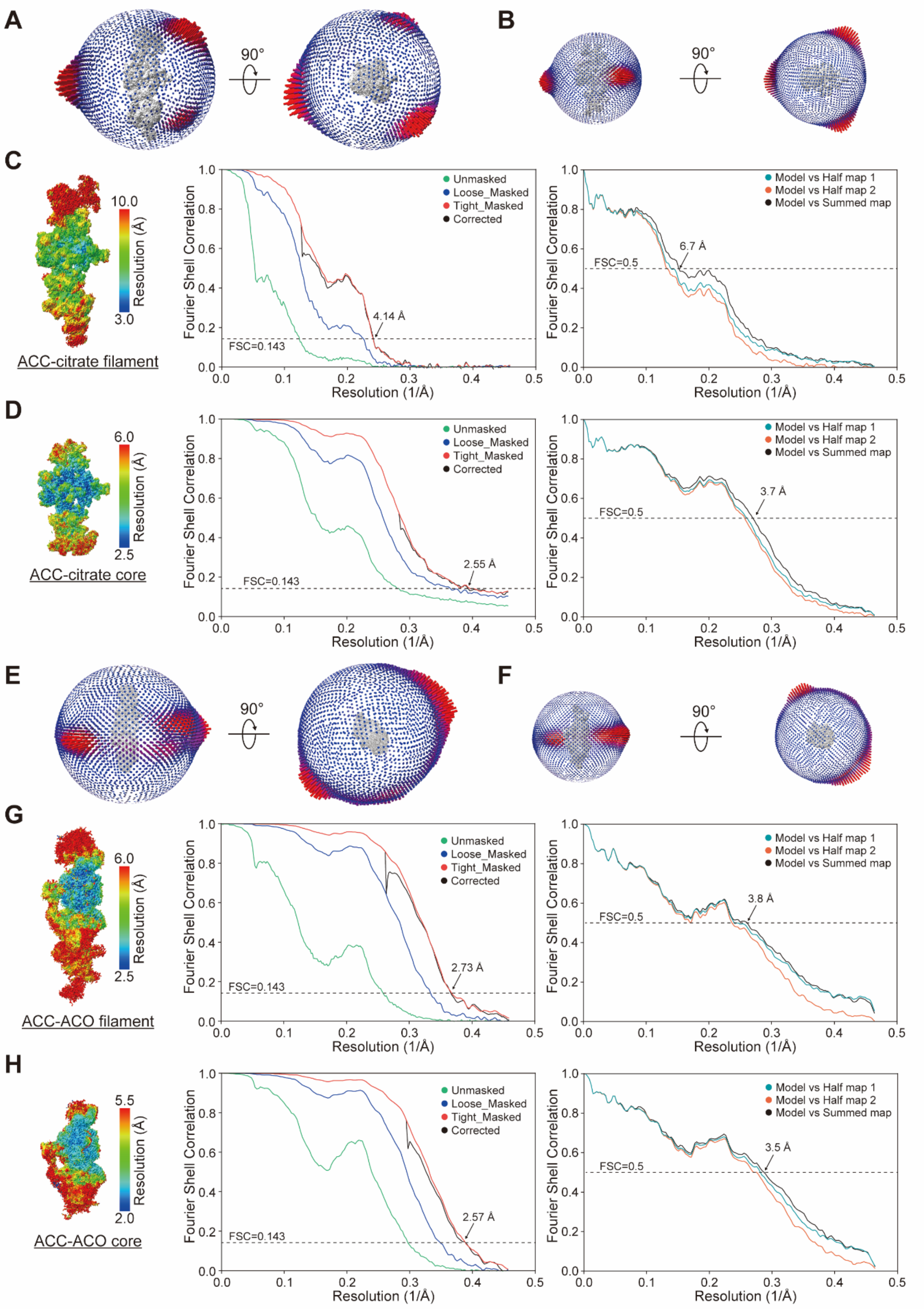
A
90°
B
90°
C
10.0
3.0
Resolution (Å)
ACC-citrate filament
Unmasked
Loose_Masked
Tight_Masked
Corrected
4.14 Å
FSC=0.143
Fourier Shell Correlation
Resolution (1/Å)
Model vs Half map 1
Model vs Half map 2
Model vs Summed map
6.7 Å
FSC=0.5
D
6.0
2.5
ACC-citrate core
2.55 Å
3.7 Å
E
90°
F
90°
G
6.0
2.5
ACC-ACO filament
2.73 Å
3.8 Å
H
5.5
2.0
ACC-ACO core
2.57 Å
3.5 Å

**Fig. S3. Cryo-EM analysis of human ACC filaments.** (**A**, **B**) Cryo-EM maps and the corresponding Euler distribution plots of the ACC-citrate filaments (**A**) and its core region (ACC-citrate core) (**B**). (**C**) Local resolution map (left), gold standard FSC curves (middle), and FSC curves for cross-validation of the structure of the ACC-citrate filaments (right). (**D**) Local resolution map (left), gold standard FSC curves (middle), and FSC curves for cross-validation of the structure of the ACC-citrate core (right). (**E**, **F**) Cryo-EM maps and the corresponding Euler distribution plots of the ACC-ACO filaments (**E**) and its core region (ACC-ACO core) (**F**). (**G**) Local resolution map (left), gold standard FSC curves (middle), and FSC curves for cross-validation of the structure model of the ACC-ACO filaments (right). (**H**) Local resolution map (left), gold standard FSC curves (middle), and FSC curves for cross-validation of the structure model of the ACC-ACO core (right). For both the ACC-citrate and ACC-ACO filaments, the particles were extracted with a box size of 488 pixels, while the particles of the core regions of the filaments were extracted with a box size of 320 pixels. The map resolution was determined by the reciprocal of the spatial frequency at FSC = 0.143, and the consistency of the structure optimization process was verified by comparison of the molecular model with the summed (the black curves) or half (the red or green curves) maps at FSC = 0.5(*28, 31*).

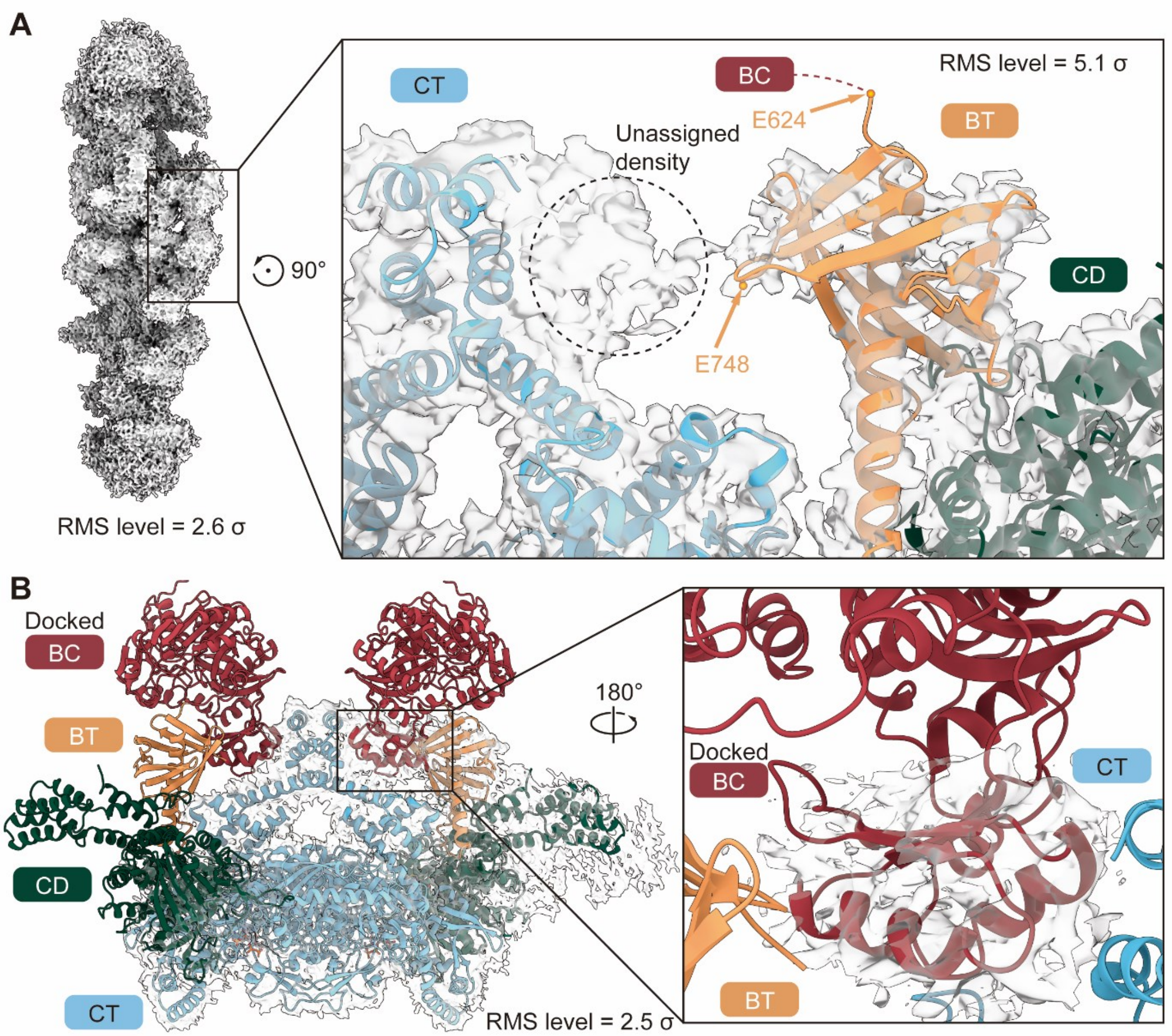


**Fig. S4. Assignment of the BC domain in the ACC-ACO filaments.** (**A**) Cryo-EM map of the ACC-ACO filaments. After building the structures of the BT, CD, and CT domains into the cryo-EM map, the unassigned density is marked by the dashed circle. The positions of the N- and C-termini (residues E624 and E748) of the BT domain are indicated by orange arrows. The map of ACC-ACO filament is displayed at RMS levels of 2.6 σ in the left panel and 5.1 σ in the right panel. (**B**) The ACC1 dimer in the structure of the ACC-ACO core structure is aligned with the corresponding cryo-EM density. Considering the positions of the BT domain and the partial density near the C-terminal of CT domains, the structure of the BC domain was docked into the unassigned density. The map of ACC-ACO core is displayed at an RMS level of 2.5 σ in the figure.

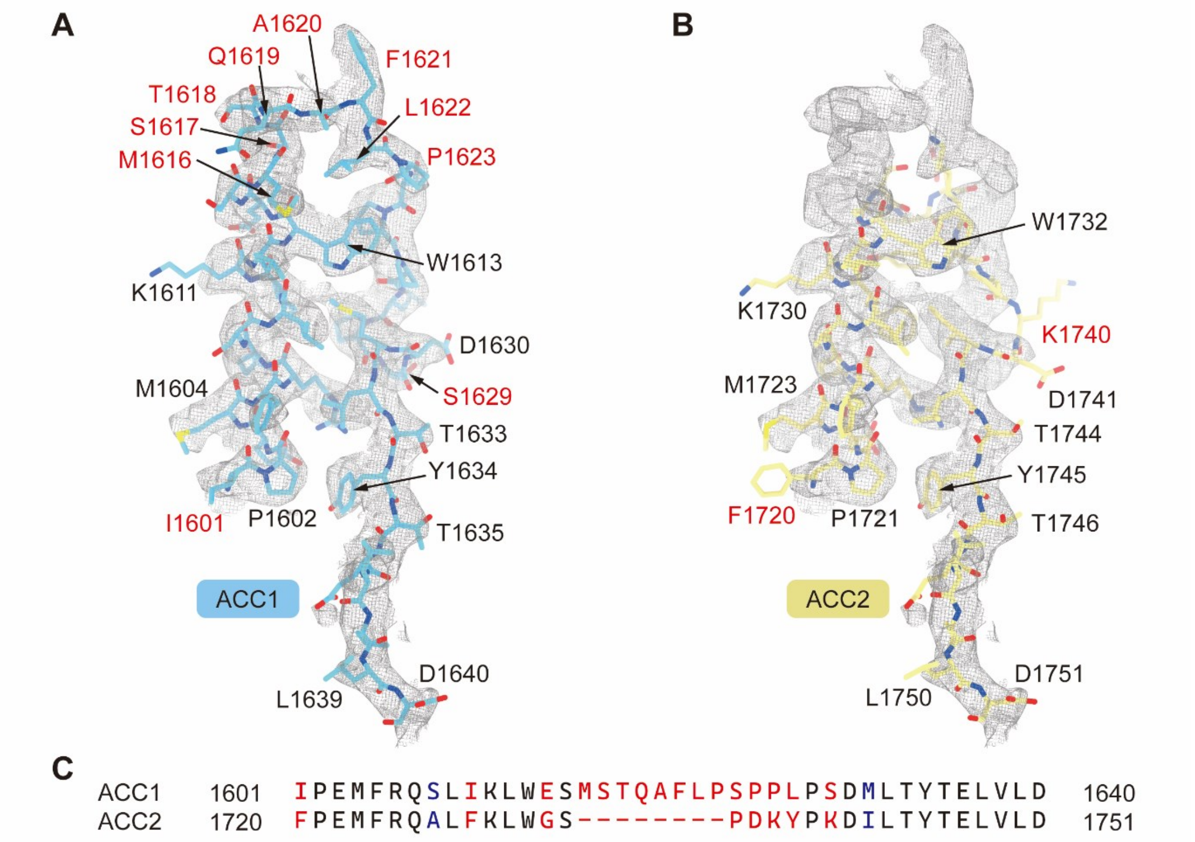


**Fig. S5. The cryo-EM map of the ACC-citrate filaments aligns with the structure of human ACC1 but not ACC2.** (**A**) Docking of the structure of human ACC1 (residues 1601–1640) into the cryo-EM map of the ACC-citrate filaments. (**B**) Docking of the structure of human ACC2 (residues 1720–1751) into the cryo-EM map of the ACC-citrate filaments. (**C**) Alignment of the protein sequence of human ACC1 (residues 1601–1640) with that of human ACC2 (residues 1720–1751). The identical, similar and not similar residues are colored black, blue and red, respectively.

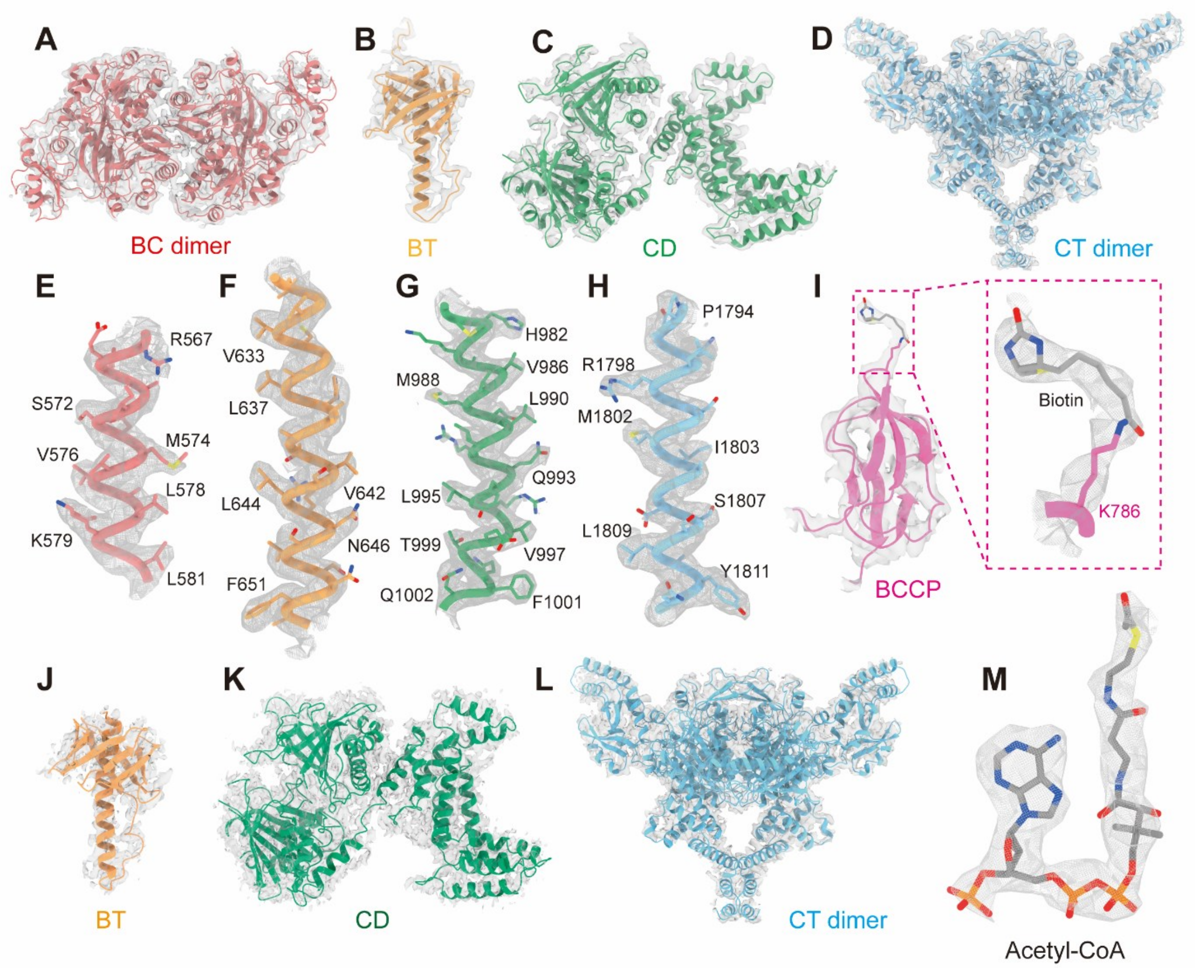


**Fig. S6. The cryo-EM maps of human ACC1 in the ACC-citrate filaments and the ACC-ACO filaments.** (**A**–**D**) Cryo-EM maps of the BC homodimer (**A**), BT domain (**B**), CD domain (**C**) and CT homodimer (**D**) in the ACC-citrate filaments. (**E**–**H**) Cryo-EM maps of representative α-helices in the BC (**E**), BT (**F**), CD (**G**) and CT (**H**) domains in the ACC-citrate filaments. (**I**) Cryo-EM map of the biotinylated BCCP domain in the ACC-citrate filaments. The side chain of K786 and the covalently linked biotin are shown as sticks. (**J**–**L**) Cyro-EM maps of the BT domain (**J**), CD domain (**K**) and CT homodimer (**L**) in the ACC-ACO filaments. (**M**) Cryo-EM map of the acetyl-CoA bound to the catalytic pocket of each CT homodimer in the ACC-ACO filaments.

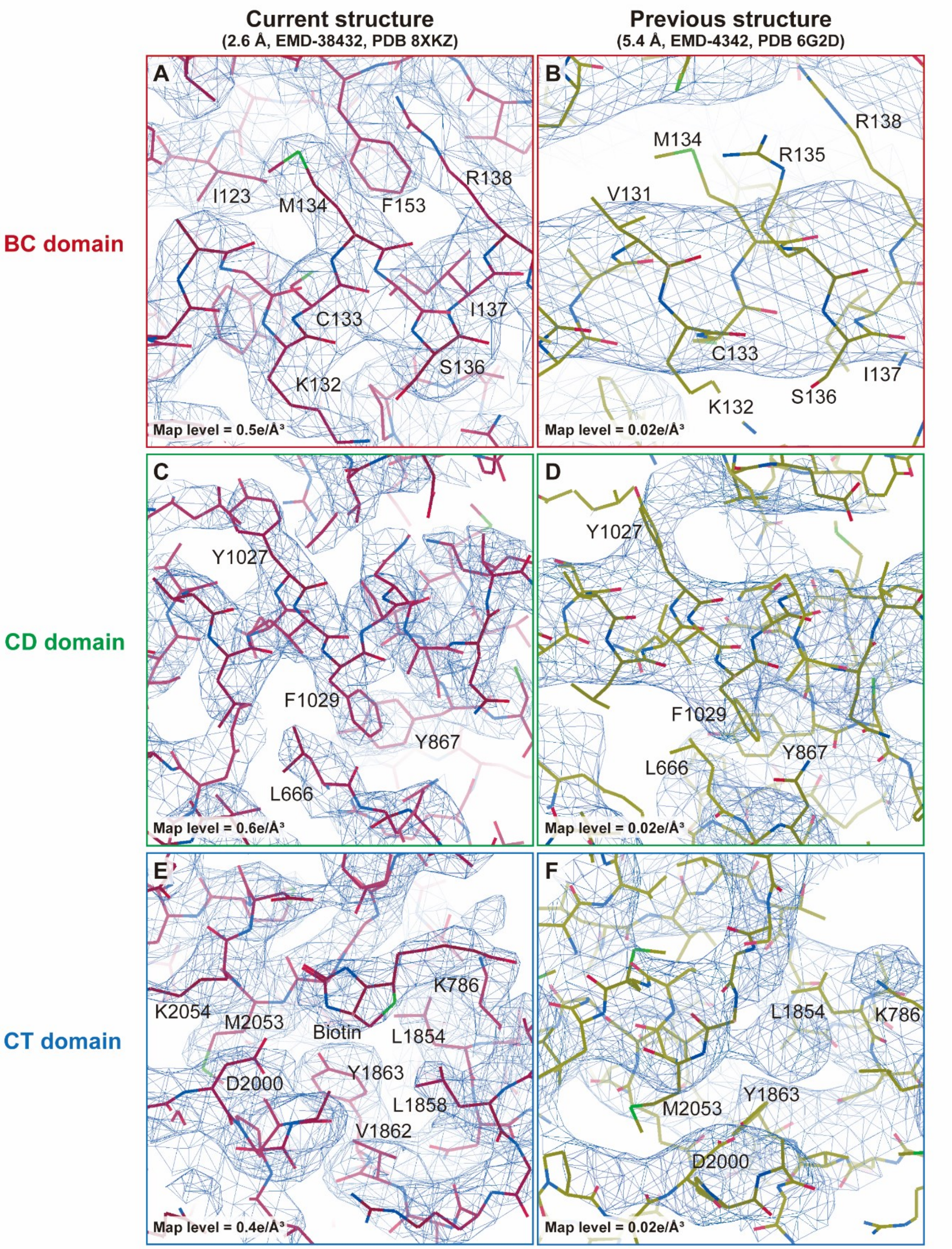


**Fig. S7. Comparison of the 2.55 Å cryo-EM map of the ACC-citrate filaments obtained in our study with the 5.4 Å cryo-EM map reported previously.** (**A**, **C**, **E**) The cryo-EM maps of the BC domain (**A**), CD domain (**C**) and the CT domain (**E**) in the structure of the ACC-citrate filaments solved in our study (PDB code: 8XKZ). (**B**, **D**, **F**) The cryo-EM maps of the BC domain (**B**), CD domain (**D**) and the CT domain (**F**) in the structure of the ACC-citrate filaments reported previously (PDB code: 6G2D)(*16*). The structures and corresponding cryo-EM maps of representative regions of the BC, BT, and CT domains in the human ACC-citrate filaments are visualized in Coot(*30*).

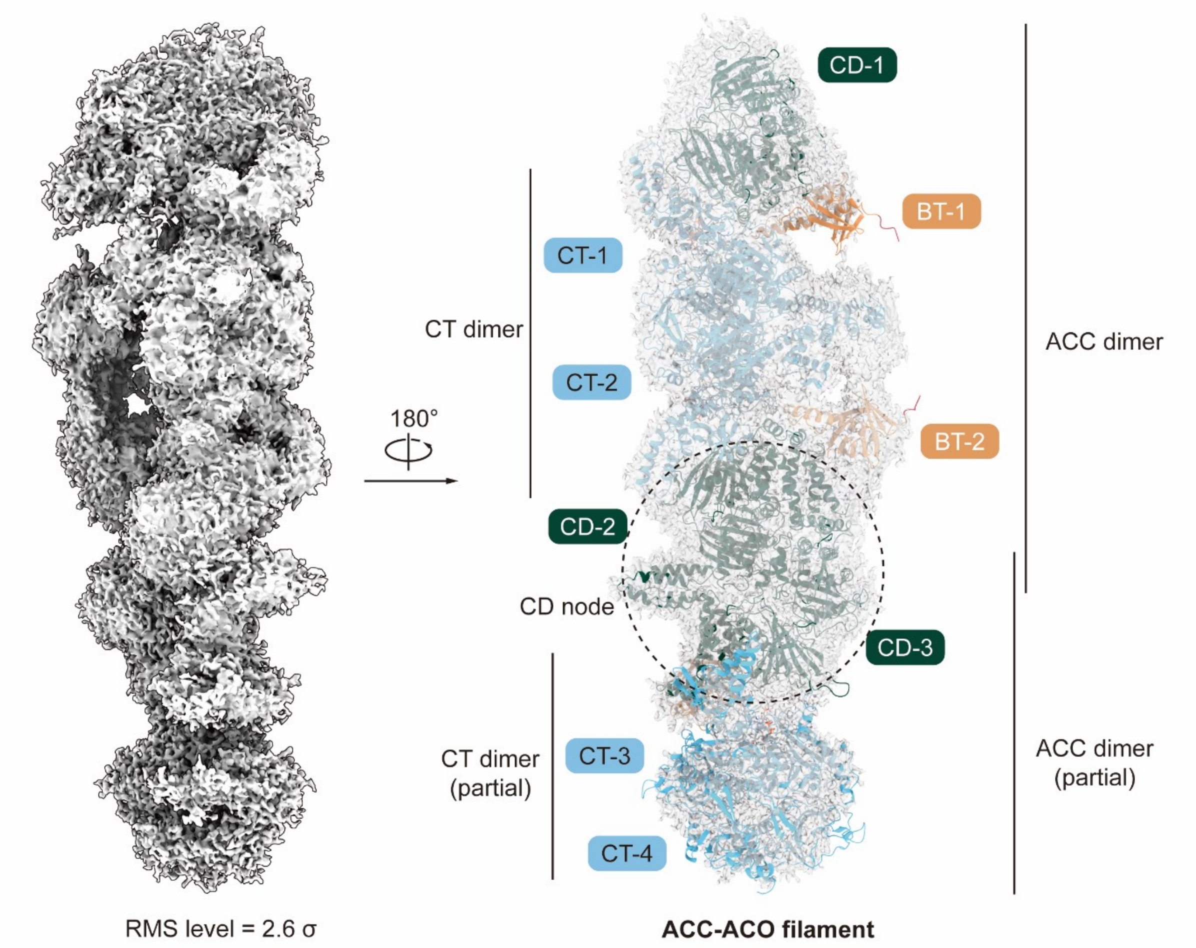


**Fig. S8. The cryo-EM map of ACC-ACO filament shows the density of the CD node.** The ACC-ACO filament structure built based on the cryo-EM map contains one and a half ACC homodimers, with the densities of the BT (orange), CD (dark green) and CT (light blue) domains recognized in the map. The CD-2 and CD-3 from adjacent ACC dimers interact with each other to form a CD node. The map of ACC-ACO filament is displayed at an RMS level of 2.6 σ.

### Supplementary Tables

**Table S1. The mass spectrometry analysis results for human ACC components.** (see the separated file *Table-S1.xlsx*)

**Table S2. Statistics of cryo-EM data collection, reconstruction, refinement, and validation.**

| **Sample** | **ACC-citrate core** | **ACC-citrate filament** | **ACC-ACO core** | **ACC-ACO filament** |
|---|---|---|---|---|
| **PDB code** | 8XKZ | 8XL0 | 8XL1 | 8XL2 |
| **EMDB code** | EMD-38432 | EMD-38433 | EMD-38434 | EMD-38435 |
| **Data collection** | | | | |
| EM equipment | Titan Krios (Thermo Fisher Scientific) | | | |
| Voltage (kV) | 300 | | | |
| Detector | Gatan K3 Summit | | | |
| Energy filter | Gatan GIF Quantum, 20 eV slit | | | |
| Pixel size (Å) | 1.0773 | | | |
| Electron dose ($e^-/Å^2$) | 50 | | | |
| Defocus range (μm) | −1.2 to −2.2 | | | |
| Number of collected micrographs | 4,669 | 4,669 | 13,632 | 13,632 |
| **3D Reconstruction** | | | | |
| Software | cryoSPARC 3.3.2 | | | |
| Number of used particles | 224,904 | 36,041 | 172,311 | 155,033 |
| Resolution (Å) | 2.55 | 4.14 | 2.57 | 2.73 |
| Symmetry | C1 | | | |
| Map sharpening B-factor ($Å^2$) | 57.2 | 85.4 | 83.2 | 81 |
| **Refinement** | | | | |
| Software | Phenix 1.14 | | | |
| ***Cell dimensions*** | | | | |
| a=b=c (Å) | 344.736 | 525.7224 | 344.736 | 525.7224 |
| $\alpha=\beta=\gamma$ (˚) | 90 | | | |
| ***Model composition*** | | | | |
| Protein residues | 4,292 | 11,692 | 2,285 | 4,901 |
| Biotin | 2 | 5 | 0 | 0 |
| Acetyl-CoA | 0 | 0 | 2 | 3 |
| ***R.m.s deviations*** | | | | |
| Bonds length (Å) | 0.011 | 0.011 | 0.009 | 0.008 |
| Bonds Angle (˚) | 0.882 | 0.884 | 0.887 | 0.822 |
| ***Ramachandran plot statistics (%)*** | | | | |
| Preferred | 93.68 | 93.85 | 96.21 | 94.76 |
| Allowed | 6.02 | 5.87 | 3.35 | 5.12 |
| Outlier | 0.30 | 0.28 | 0.44 | 0.12 |
